# The recognition and expectations of ex-inpatients of mental health services: A web-based questionnaire survey in Japan

**DOI:** 10.1101/317057

**Authors:** Akihiro Shiina, Yasutaka Ojio, Aiko Sato, Naoya Sugiyama, Masaomi Iyo, Chiyo Fujii

**Affiliations:** Division of Medical Treatment and Rehabilitation, Chiba University Center for Forensic Mental Health, Chiba-shi, Chiba, Japan; National Institute of Mental Health, National Center of Neurology and Psychiatry, Kodaira-shi, Tokyo, Japan; Department of Psychiatry, Chiba University Hospital, Chiba-shi, Chiba, Japan; Numazu Chuo Hospital, Numazu-shi, Shizuoka, Japan

**Author notes:** These authors contributed equally to this work.

## Abstract

Concern about mental health issues and the treatment of mentally disordered offenders attracts a lot of public attention. This study aimed to gather the experiences and opinions of people who have experienced admission to a psychiatric ward in order to understand their reaction to, and understanding of, the legislation behind the involuntary admission of psychiatric patients. A web-based questionnaire survey was conducted with a total of 379 participants, using a cross-sectional, exploratory design. The data were analyzed using a chi-square test, Fisher’s exact test, and a logistic regression analysis. According to the results, many patients were satisfied with their psychiatric admission, however, few participants said that they had been given adequate explanation for their involuntary treatment. Most participants expected qualified aids after discharge, although the prospect of a regular visit from an official was not entirely supported by the participants. Patient satisfaction was relevant to the discussion of their needs after discharge and in developing a crisis plan during admission. These findings suggests that psychiatric patients accept inpatient treatment as long as they receive an adequate explanation. More qualified care such as relapse prevention would be expected to lead to better satisfaction. In order for them to welcome regular visits from an official, patients may need more information and discussion.

## Introduction

In recent years, mental health has become one of the most devastating health concerns faced both by mental health professionals and the general public. The World Health Organization (WHO) considers several mental disorders to be a cause of living through a damaging disability-adjusted life year (DALY) [1]. In many nations, especially in developed countries, public attention has been caught by the rise of mental disorders, and in finding effective treatments for them.

In addition, mental disorders are often focused on as a cause of criminal offenses, in public perception and in the media. There have been several cases in which the perpetrator of a serious crime has been found to have been suffering from a mental disorder. These incidents often attract public anger towards, and suspicion of, offenders with mental disorders. This public opinion has led to a discussion of the need for a new provision within the legal system for treating offenders with mental health issues in each nation [2-5].

In Japan, in 2016, a massacre occurred at a residence for disabled people in Sagamihara city. The defendant, who was an ex-employee of the residence, was suspected to have broken into the facility to kill 19 residents as a result of his prejudiced ideology [6]. He had been hospitalized involuntarily for a couple of weeks by order of the prefectural governor under the Mental Health and Welfare Act (MHWA), and was likely an abuser of cannabis. This incident ignited a broad public argument about several issues relevant to the current situation in Japan. Following the report published by the special team in charge of examining the incident [7], the government submitted an amendment of the MHWA. The amended bill contained a new scheme for the official follow-up of patients who were hospitalized by the prefectural governor’s order. Some politicians, as well as scholars, disagreed with the bill, as they were concerned about the potential risk that the patients under supervision would experience a restriction of their human rights. The bill was abandoned because of the dissolution of the Lower House due to another political conflict.

The corresponding author has been a member of a research team focused on reforming the MHWA. Through a number of team conferences, the need for a survey that gathers the opinion of the subjects of this legislation (i.e. offenders with mental disorders) was suggested. Gaining an understanding of how the people who have experienced admission to a psychiatric ward feel about the current situation of mental health care, and what they want for the future, was considered to be essential before discussing the validity of the new scheme suggested by the bill. On the basis of these recommendations, the present study aimed to clarify the opinion of psychiatric patients, who have had experiences of admission, about the mental health care for inpatients. It also aimed to understand whether they want to receive the mental health care that it is estimated would be provided based on the predicted reforms of the MHWA. This study was conducted using a cross-sectional, exploratory, web-based questionnaire survey.

## Materials and methods

### Materials

We developed a series of questions based on a discussion conducted by the research group members. The items from the questionnaire are shown in the appendix. Only people who had experience of admission to a psychiatric ward participated in the survey. There were five sections in the questionnaire. In Section One, participants were asked about their knowledge and opinion of the MHWA and the Medical Treatment and Supervision Act (MTSA), which was created in 2005 for the treatment of those who had committed serious harm due to psychiatric conditions. The content of these questions was based on those from a previous study in which the corresponding author asked for the knowledge and opinions about forensic mental health from outpatients with a mental disorder [8]. In Section Two, participants were asked about the form of admission they had experienced (either MHWA or MTSA). In Section Three, participants were asked about their opinion of their latest psychiatric admission. In Section Four, participants were presented with examples of treatment for aiding inpatients in preparation of discharge and in the prevention of readmission. Participants were asked whether each of these examples was offered to them during their latest psychiatric admission. Finally, in Section Five, we asked whether, if they were to be readmitted to a psychiatric ward, the participants wanted to receive the types of aid that had been presented to them in the previous section.

### Procedure

The survey was conducted from April 27th to May 31^st^, 2017. We made a contract with the Japan Research Center (JRC), a marketing company in Japan, for the implementation of the survey. Cyber Panel was used (http://www.nrc.co.jp/monitor/cyber201.html), which is a registration system of web-based questionnaires managed by JRC, to search the database consisting of approximately 200,000 people. JRC sent the questionnaire to any of those identified in the database with a medical history of any mental disorders. The respondents were informed of the aim of this study, that personal information of the participants would not be sent to us, that it would cost them nothing (except telecommunication costs), that the results would be published, and that participants would be rewarded by JRC.

### Data analysis

After completion of the survey, the anonymized data were sent to us. We analyzed the data using IBM SPSS Statistics for Windows, Version 24 (IBM Corp., Armonk, NY, United States), and set the level of significance *at p* <0.05.

### Ethical issues

The study protocol was approved by the ethics committee of the Graduate School of Medicine at Chiba University on April 27^th^, 2017 (no. 189). We did not receive any personal information from the participants or JRC. All respondents agreed to participate by sending in their answer form. We registered the study with the Clinical Trials Registry of the University Hospital Medical Information Network (UMIN, Tokyo, Japan) with the unique trial number UMIN000027316.

## Results and discussion

### Demographic data

JRC identified a total of 35,505 people with mental disorders as candidates for the survey. The categories of disorders were: depression (9,644), sleep disorders (6,082), neuroses (5,189), panic disorder (3,404), bipolar disorder (1,672), bulimia nervosa (1,579), social anxiety disorder (1,378), anorexia nervosa (1,222), obsessive-compulsive disorder (1,005), schizophrenia (925), post-traumatic stress disorder (892), general anxiety disorder (815), attention-deficit hyperactivity disorder (446), and other mental disorders (1,252). JRC sent the request form of the questionnaire to all 35,505 registrars. A total of 379 participants who had at least one experience of admission to a psychiatric ward replied to the questionnaire. All of their data were included in the following analyses.

### Screening

The departments of medical facilities the participants had consulted were: psychiatry (379/100%), internal medicine (358/94.5%), dentistry (347/91.6%), surgery (256/67.5%), pediatrics (158/41.7%), gynecology (136/35.9%), and other departments (169/44.6%). The departments of medical facilities the participants had been hospitalized in were: psychiatry (379/100%), internal medicine (135/35.6%), surgery (116/30.6%), gynecology (72/19.0%), pediatrics (16/4.2%), dentistry (12/3.2%), and other departments (52/13.7%).

### Section One

A total of 50 (13.2%) participants answered that they knew MHLA well, and a total of 79 (20.8%) knew that the MTSA had been enforced. The proportion who answered that they knew the MHWA well was higher than that of a previous survey conducted by the corresponding author using psychiatric outpatients (Chi-square test, df=2, Chi-square=197.43, *p*<0.001. See fig 1) [8]. With regards to the scheme of involuntary admission by the prefectural governor’s order, 149 (39.3%) were definitely affirmative of the policy and 110 (29%) were relatively affirmative, whereas 4 (1.1%) were definitely against and 10 (2.6%) were relatively against. With regards to the scheme of MTSA, 159 (42%) were affirmative and 92 (24.3%) were relatively affirmative, whereas 3 (0.8%) were definitely against and 12 (3.2%) were relatively against.

**Fig 1.**
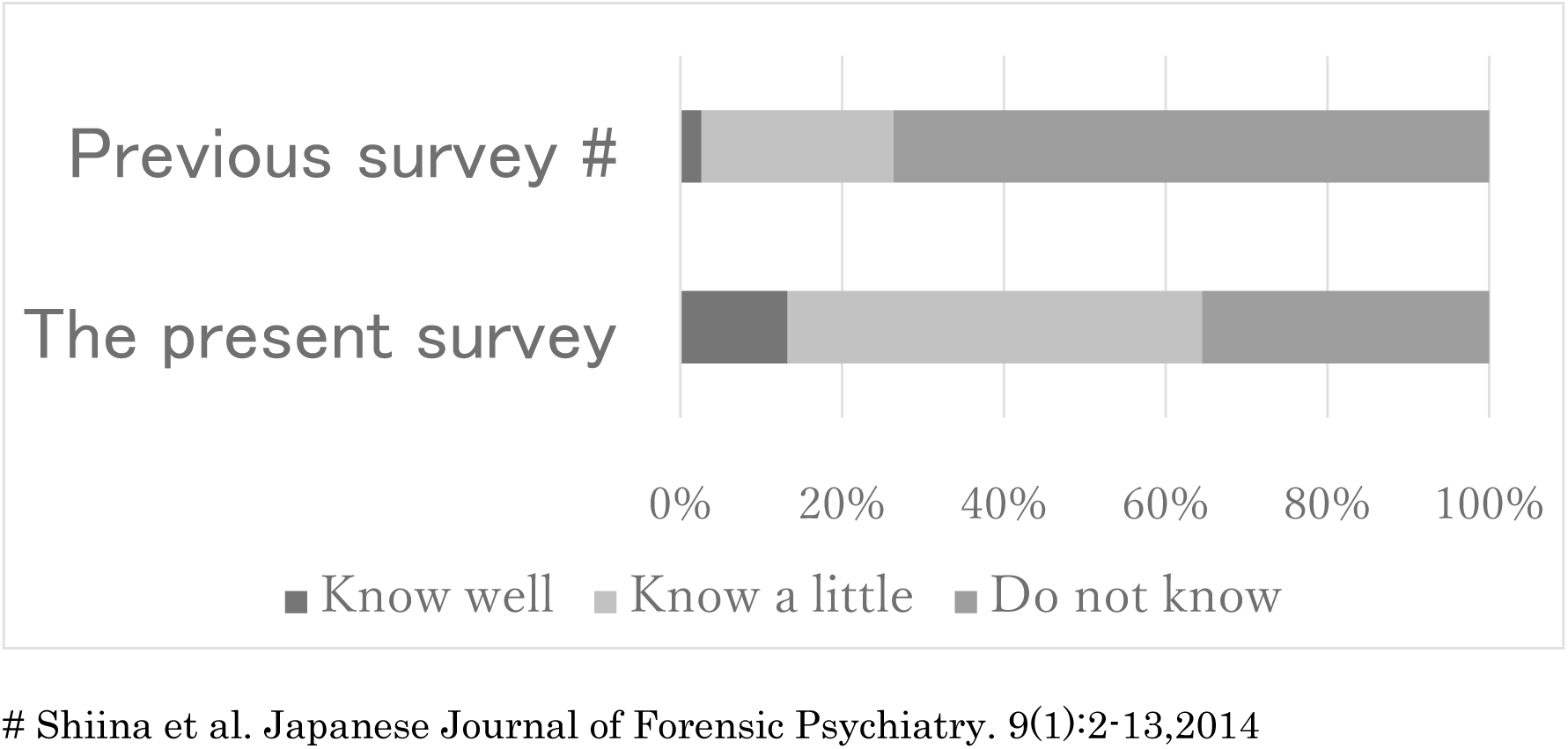
Knowledge regarding the Mental Health and Welfare Act (compared with a previous survey)

### Section Two

In total, 45 (11.9%) participants experienced involuntary admission by the prefectural governor’s order in relation to the MHWA (including urgent involuntary admission, limited up to 72 hours). In addition, 35 (9.2%) experienced admission for medical care and protection through the MHWA, 98 (25.9%) experienced voluntary admission through the MHWA, 11 (2.9%) experienced emergency admission through the MHWA, 7 (1.8%) experienced hospitalization order by the court through the MTSA, and 4 (1.1%) experienced hospitalization for assessment through the MTSA. A total of 102 (26.9%) participants answered that they experienced another form of admission to a psychiatric ward, and 102 (26.9%) did not know the form of admission they had experienced.

These findings were analyzed by cross-tabulation, which revealed that the participants who experienced involuntary admission by the prefectural governor’s order had a significantly more unfavorable opinion to this scheme than those without the experience of involuntary admission (Chi-square test, df=2, Chi-square=9.004, *p*=0.011. See Fig 2).

**Fig 2.**
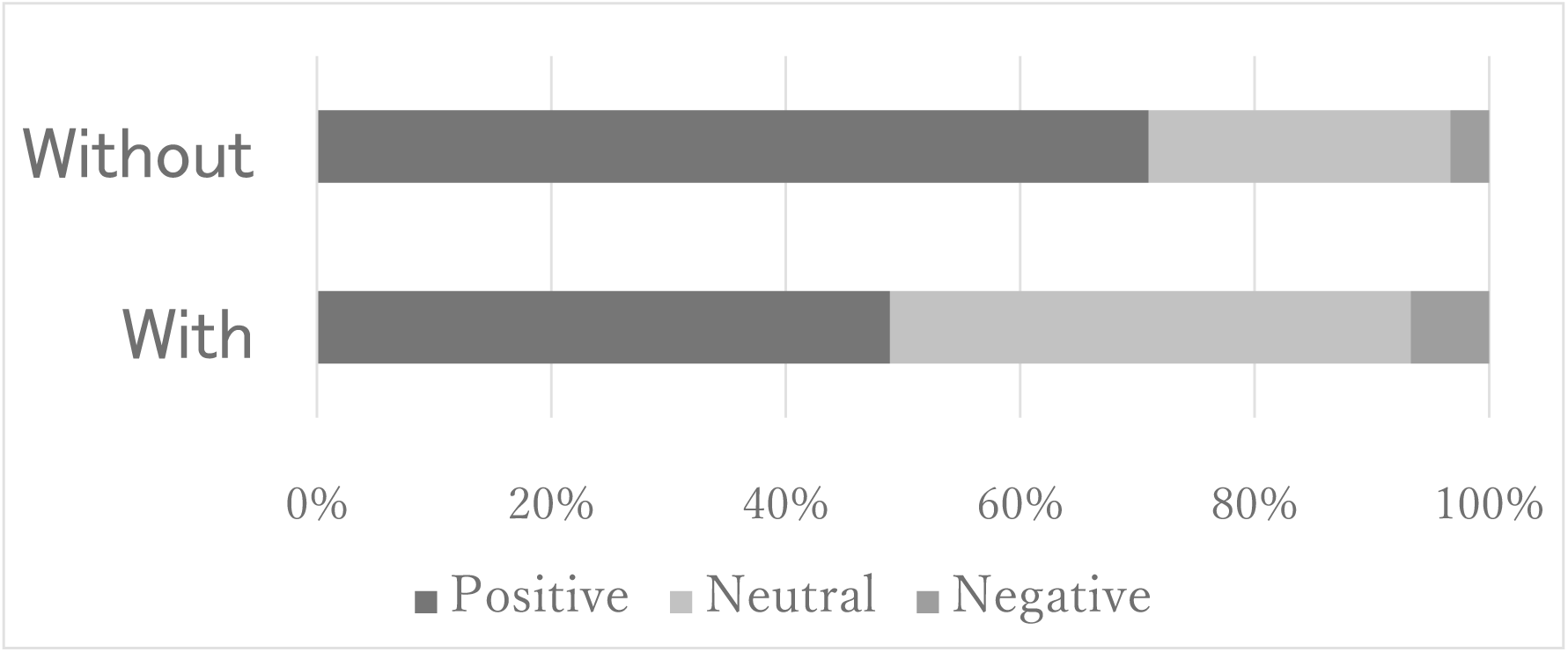
Opinion toward involuntary admission by the prefectural governor’s order (Split by whether or not involuntary admission by the prefectural governor’s order was experienced)

### Section Three

The form of latest admission to a psychiatric ward experienced by the participants were: involuntary admission by the prefectural governor’s order in the MHWA (including urgent involuntary admission) (30/7.9%), admission for medical care and protection in the MHWA (19/5%), voluntary admission in the MHWA (88/23.2%), emergency admission in the MHWA (6/1.6%), hospitalization order by the court in the MTSA (5/1.3%) hospitalization for assessment in the MTSA (3/0.8%), other forms of admission (126/33.2%), respectively. A total of 102 (26.9%) participants did not know the form of their latest admission.

The majority of the participants answered that they had accepted the explanation of the necessity of admission. Approximately one-third of the participants felt themselves at risk of harm self or others at the time of the latest psychiatric admission. Almost one-fifth of the participants received treatment without their consent. Nearly two-thirds of the participants believed that their latest admission was necessary. Limited to those whose form of admission was an involuntary admission by the prefectural governor’s order, 19 (67.9%) felt the admission had been necessary. Almost half of the participants answered that they were definitely or relatively satisfied with the treatment they received during their latest admission. These results are shown at the table 1.

**Table 1.**
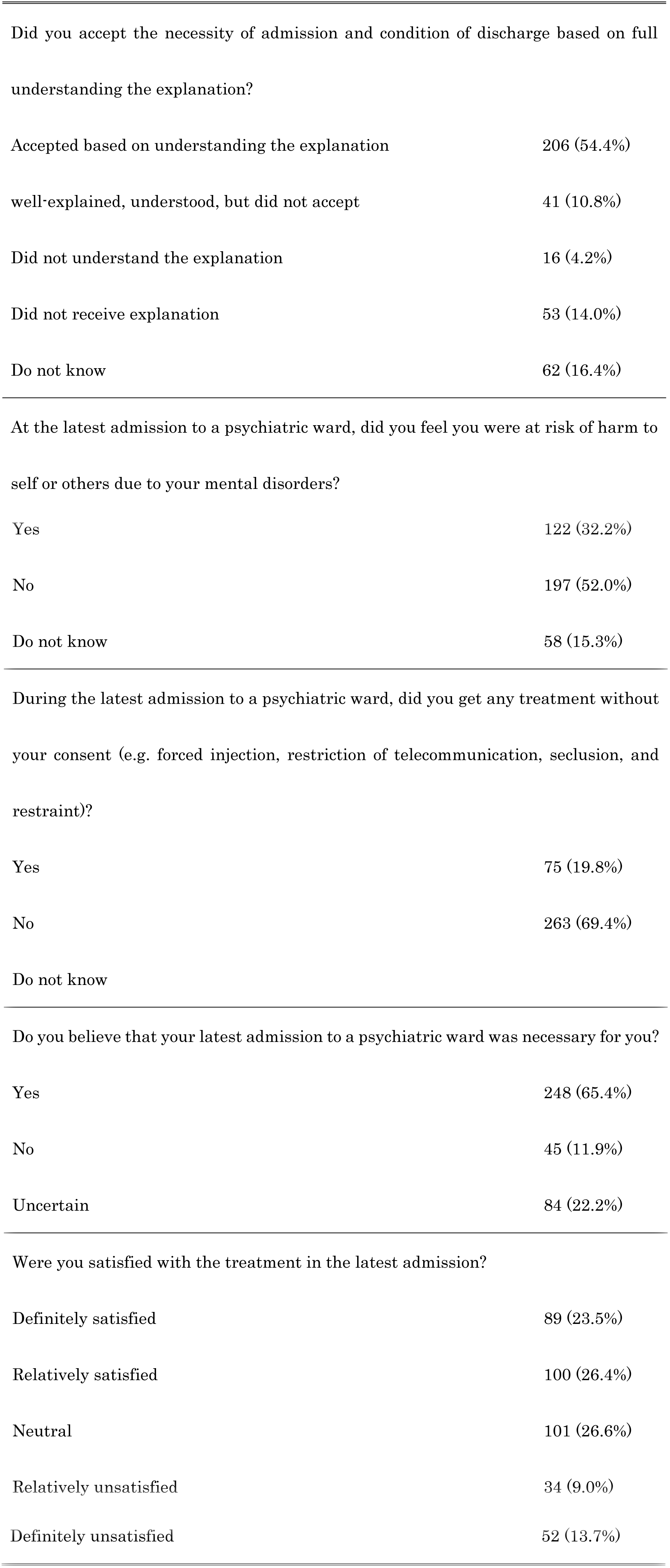
Recognition of their latest psychiatric admission

### Section Four

In all ten of the items, a minority of the participants answered that they had been given the types of aids that were described (See Fig 3). Only 40 (10.1%) of the participants answered that they had been given a clear explanation about the necessity of involuntary treatment. The proportion was not significantly different between those who actually received treatment without their consent and those did not receive any involuntary treatments (See Fig 4).

**Fig 3.**
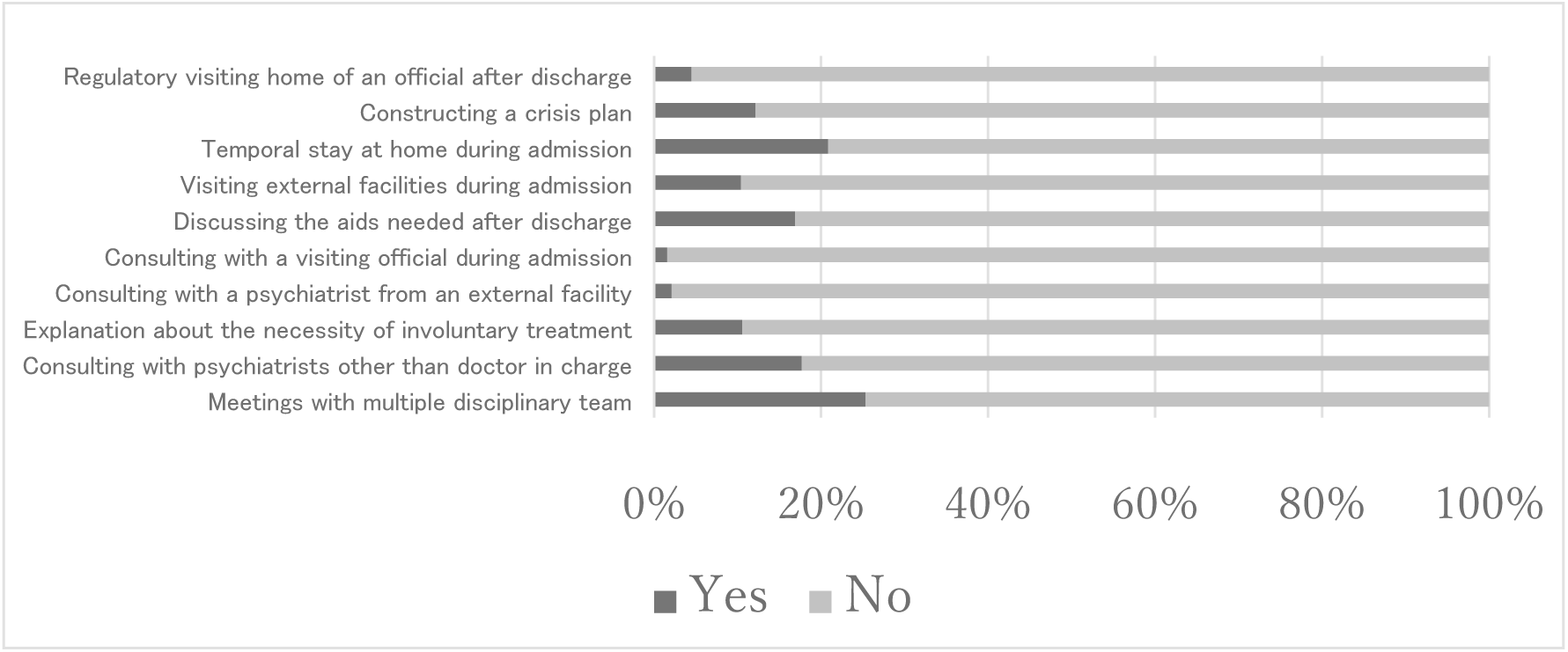
Aids given during the latest admission

**Fig 4.**
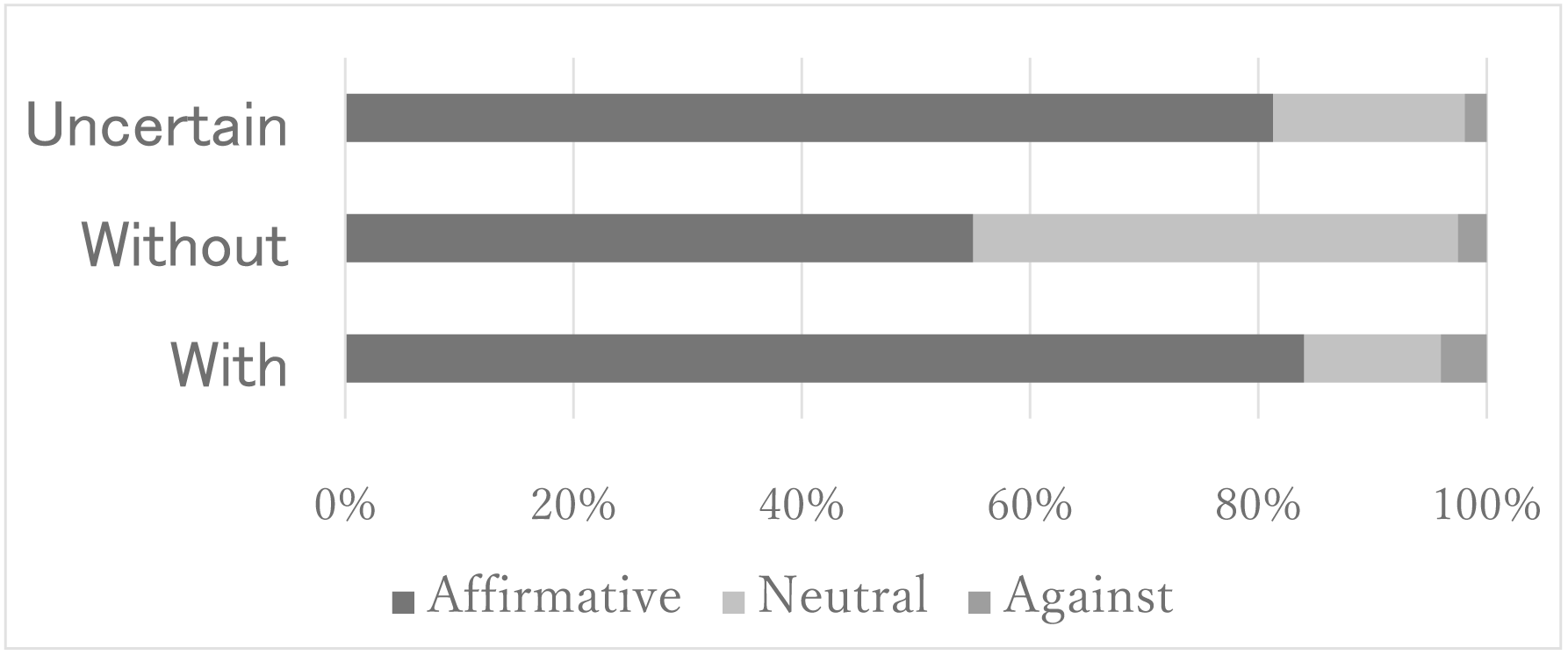
Opinion toward receiving clear explanation about the necessity of involuntary treatment (split by whether coercive treatment was experienced)

Also, we converted the answers of the latest psychiatric admission satisfaction level into a binary scale: “definitely satisfied” and “relatively satisfied” were converted into “satisfactory,” and “neutral”, “relatively unsatisfied,” and “definitely unsatisfied” were converted into “unsatisfactory.” After the conversion, 189 (49.4%) participants were classified as “satisfactory,” while 187 (49.3%) were classified as “unsatisfactory.”

Next, we examined whether their satisfaction was altered according to the explanation about the necessity of involuntary treatment. With Fisher’s exact test, we found no significant difference in the proportion of satisfied participants, regardless of the explanation given (Chi-square test, P=0.091). However, when stratification with the existence of involuntary treatment was applied, among the participants who received involuntary treatments, those who had received an explanation of the necessity of involuntary treatment were significantly more likely to be satisfied than those who had not (Fisher’s exact test, *p*=0.016.)

Furthermore, we applied a logistic regression analysis to identify the factors that influenced the satisfaction of the inpatients. We set the binary value “satisfaction” above as the dependent variable and input the ten items of treatment for aiding inpatients (from Section 4) as the independent variables. We used stepwise, logistic regression analysis, with increasing variables. As a result, two items: “discussing the aids needed after discharge (B=0.886, SE=0.317, Wald=7.819, df=1, *p*<0.005, Exp(B)=2.427)” and “constructing a crisis plan (B=0.847, SE=0.375, Wald=5.107, df=1, *p*<0.024, Exp(B)=2.322)” were extracted as significantly relevant to satisfaction level.

### Section Five

In all of the ten items, positive opinions were much more prevalent than negative ones. However, with the items “consulting with a visiting official (e.g., public health nurse) during admission” and “regulatory visiting home of an official (e.g., public health nurse) after discharge,” a considerable number of participants answered that they were uncertain (See Fig 5).

**Fig 5.**
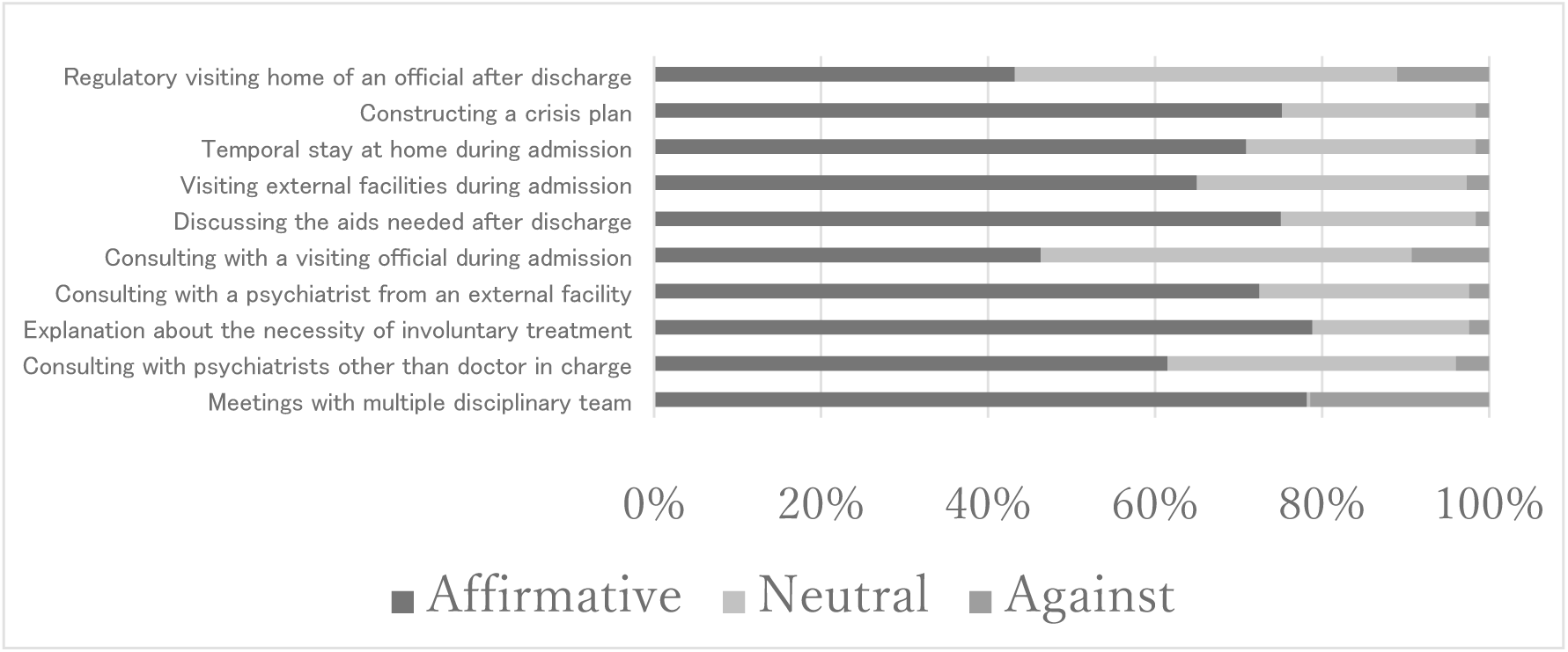
Opinions about the aids in a future admission

## Discussion

In the present study, we conducted a questionnaire with people who had experience of admissions to a psychiatric ward. This survey focused on patients’ opinion toward mental health care services. The results from this study give us some important information regarding the opinions of psychiatric patients. Over 35,000 patients (approximately one percent of all psychiatric patients in Japan) [9] were screened to detect more than 300 participants with experiences of psychiatric admission. We are aware of no surveys of a similar scale using psychiatric patients in Japan.

In this study, a considerable proportion of the participants answered that they had knowledge of the legislation regarding mental health services. Ex-inpatients are more likely to know the MHWA well, compared to general outpatients. These findings suggest that individuals are best able to gain knowledge of the mental health law through their own experience. On the other hand, it appears that the participants did not possess enough knowledge of their psychiatric admission. A total of 126 (33.2%) participants answered that their latest form of admission to a psychiatric ward was “other forms of admission.” It is, however, extremely rare for a patient to be hospitalized neither under MHWA nor MTSA [10]. Thus, it seems that up to a third of the participants may have misunderstood the form of admission they experienced, and so it is possible that some patients are not as familiar with mental health law as much as they believe. Alternatively, some patients may have not been concerned about their legal status at the time of their admission, so may not have paid attention to the information provided to them.

Many participants admitted at the time of the survey that their latest admission had been necessary. This result is similar to those in studies previously conducted in other nations in which Switzerland [11] and Ireland [12]. However, the participants whose latest admission was an involuntary admission by the prefectural governor’s order had a relatively negative opinion toward involuntary admission (see fig 2.) This result is consistent with that of a previous study in Japan [8]. It means that the scheme of involuntary admission by the prefectural governor’s order may require the fostering of a better level of understanding in its patients in order to increase the level of satisfaction.

An additional analysis revealed that the satisfaction of the participants who received involuntary treatments was associated with a thorough explanation. The high satisfaction level of patients may motivate them to continue treatment, leading to good outcome [13]. Thus, in cases where patients are confused about the necessity of the treatment, health practitioners should spend more time explaining it.

Although we believed before conducting the survey that many psychiatric hospitals had already been offering aids for a qualified care, most participants answered that they had not been given them. The distribution of each answer was not altered by sorting by the form of admission. Among them, the result that only 10.1% of the participants believed that they received a proper explanation regarding the necessity of involuntary treatment is surprising. The MHWA requires practitioners to explain the possibility of involuntary treatment to all inpatients at the beginning of admission. In reality, it appears that most patients did not understand the explanation that was provided to them. If there are cases where medical practitioners adopt coercive means to provide treatment without proper explanation to psychiatric inpatients, this is something that needs to be addressed as a matter of priority.

On the other hand, positive opinions were more common than negative ones in every item which were asked in the section 5. The participants overall seemed happy to be given the aids which were planned to be introduced after the amendment of the MHWA. Patient satisfaction was positively associated with discussing the aids needed after discharge and with constructing a crisis plan. These items are interpreted as preventing readmission. Identifying the risk of readmission with providing some solutions in advance to the situation are important, not only for good treatment outcomes but also for better patient satisfaction.

An exception to this is that visiting officials to patient’s home were not entirely welcomed. Not many participants actively opposed the proposal, but half withheld their opinion towards receiving an official’s visit after being discharged. It is possible that patients are not willing to be involved in a scheme of official support for discharged patients. Or perhaps they have an inaccurate perception of what the official visit would entail.

A major limitation of this study is the sampling bias. The participants were people who had at least one experience of psychiatric admission. Thus, it is possible that their mental disorders had been more severe than those in general psychiatric patients. On the other hand, the participants had registered voluntarily to JRC, which suggests that they were capable of making an account of internet service and were willing to respond to some social surveys. Thus, the participants of this study were those recovered from a relatively severe mental state. Overall, it may not be the case that the participants are representative of psychiatric patients as a whole. Regardless, the results from this study did provide important information regarding the opinions of psychiatric patients, due to the large screening population and sample size [9].

## Conclusions

We conducted a national web-based anonymous questionnaire survey to gather data from hundreds of participants with experience of psychiatric admission. Although the majority of them accepted the necessity of involuntary admission, they felt they were not given a proper explanation about it at the latest admission. The contents of support planned in amended MHWA seem to be welcomed by patients. The best way of dealing with an official visiting their home after a patient’s discharge should be discussed more concretely.

## Acknowledgement

This study was supported by a grant to the corresponding author from the Japanese Ministry of Health, Labour and Welfare as part of a research project entitled “Research of the inclusive care for the psychiatric patients discharged from involuntary admission by the prefectural governor’s order.” The authors declare no potential conflicts of interest concerning the research, authorship, and publication of this article.

Partial results from this study were presented at the 37th annual meeting of the Japanese Society for Social Psychiatry (on March 2nd, 2017, Osaka, Japan).

## Appendix: The items of the questionnaire

## Section 1

(1) Do you know the contents of the Mental Health and Welfare Act? [know well/ know a little/ do not know]
(2) Do you know the enforcement of the Medical Treatment and Supervision Act in 2005? [Know/ Do not know]
(3) What is your opinion toward the scheme of involuntary hospitalization by the prefectural governor’s order for the patients at risk of harm self or others due to mental disorders? [Definitely agree/ Relatively agree/ Neutral/ Relatively disagree/ Definitely disagree]
(4) What is your opinion toward the scheme of the Medical Treatment and Supervision Act for preventing relapse and reintegration to the society of patients who had committed serious harms due to mental disorders? [Definitely agree/ Relatively agree/ Neutral/ Relatively disagree/ Definitely disagree]

## Section 2

Choose all the forms of admission you have experienced in a psychiatric ward (multiple choice admitted). [Involuntary admission by the prefectural governor’s order in the MHWA, admission for medical care and protection in the MHWA, voluntary admission in the MHWA, emergency admission in the MHWA, hospitalization order by the court in the MTSA, hospitalization for assessment in the MTSA, other form of admission to a psychiatric ward, or form unknown]

## Section 3

(1) What is the latest form of admission to a psychiatric ward that you experienced? [Involuntary admission by the prefectural governor’s order in the MHWA, admission for medical care and protection in the MHWA, voluntary admission in the MHWA, emergency admission in the MHWA, hospitalization order by the court in the MTSA, hospitalization for assessment in the MTSA, other form of admission to a psychiatric ward, or form unknown]
(2) At the latest admission to a psychiatric ward, did you accept the necessity of admission and condition of discharge based on full understanding the explanation? [Accepted based on understanding the explanation; well-explained, understood, but did not accept; did not understand the explanation; did not receive explanation; and do not know]
(3) At the latest admission to a psychiatric ward, did you feel you were at risk of harm to self or others due to your mental disorders? [Yes/ No/ Do not know]
(4) During the latest admission to a psychiatric ward, did you get any treatment without your consent (e.g. forced injection, restriction of telecommunication, seclusion, and restraint)? [Yes/ No/ Do not know]
(5) Do you believe that your latest admission to a psychiatric ward was necessary for you? [Yes/ No/ Uncertain]
(6) Were you satisfied with the treatment in the latest admission? [Definitely satisfied/ Relatively satisfied/ Neutral/ Relatively unsatisfied /Definitely unsatisfied]

## Section 4

Choose the contents you received during the latest admission to a psychiatric ward (multiple choice admitted).

(1) Meetings with multiple disciplinary teams
(2) Consulting with psychiatrists other than the doctor in charge
(3) Receiving clear explanations about the necessity of involuntary treatment
(4) Consulting with a psychiatrist from an external facility when you do not accept the inpatient treatment
(5) Consulting with a visiting official (e.g. public health nurse) during admission
(6) Discussing the contents of aids estimated to be needed after discharge
(7) Visiting external facilities which would be concerned after discharge (e.g. day care center) during admission
(8) Temporal stay at home during admission
(9) Constructing a crisis plan
(10) Regular visits to the home of an official (e.g. public health nurse) after discharge

## Section 5

Do you agree with each of the ideas below if you have to readmit to a psychiatric ward? [Yes/ No/ Uncertain]

